# Hitchhiking, collapse and contingency in phage infections of migrating bacterial populations

**DOI:** 10.1101/378596

**Authors:** Derek Ping, Tong Wang, David T. Fraebel, Sergei Maslov, Kim Sneppen, Seppe Kuehn

## Abstract

Natural bacterial populations are subject to constant predation pressure by phages. Bacteria use a variety of well-studied molecular mechanisms to defend themselves from phage predation. However, since phage are non-motile, perhaps the simplest defense against phage would be for bacteria to outrun their predators. In particular, chemotaxis, the active migration of bacteria up attractant gradients, may help the bacteria escape slowly diffusing phages. Here we study phage infection dynamics in migrating bacterial populations driven by chemotaxis through low viscosity agar plates. We find that expanding phage-bacteria populations support two migrating fronts, an outermost “bacterial” front driven by nutrient uptake and chemotaxis and an inner “phage” front at which bacterial population collapses due to phage predation. We show that with increasing adsorption rate and initial phage population, the rate of migration of the phage front increases, eventually overtaking the bacterial front and driving the system across a “phage transition” from a regime where bacteria outrun a phage infection to one where they must evolve phage resistance to survive. We confirm experimentally that this process requires phages to “surf” the bacterial front by repeatedly reinfecting the fastest moving bacteria. A deterministic model recapitulates the transition. Macroscopic fluctuations in bacterial densities at the phage front suggest that a feedback mechanism, possibly due to growth rate dependent phage infection rates, drives millimeter scale spatial structure in phage-bacteria populations. Our work opens a new, spatiotemporal, line of investigation into the eco-evolutionary struggle between bacteria and their phage predators.

**Significance Statement:** The infection of bacteria by phage requires physical contact. This fact means that motile bacteria may avoid non-motile phage by simply running away. By this mechanism bacterial chemotaxis may help bacteria to escape phages. Here we show that when phage infect bacteria moving in soft agar plates, high phage populations or infectivity rates result in phages stopping and killing all bacteria. Conversely, when initial phage numbers or infectivity rates are low, bacteria are able to migrate away from phage successfully, despite phage ability to “surf” bacterial fronts for more than 24 hours. Between these regimes we document a “phage transition” where bacterial physiology and contingency in phage infection manifest through large-scale fluctuations in spatio-temporal dynamics.

## Introduction

Although most real-world habitats of phages and their bacterial hosts have structure at multiple spatial scales,our quantitative understanding of the dynamics of phage infections is still largely based on experiments in well-mixed systems and mass action models (1–6). Several attempts to describe spatially-structured phage-bacterial interactions have been made recently (7–9), including the experiments that quantify plaque morphology (10), colony growth (11), and the persistent stage of phage infections (12).

Here, for the first time, we study the interaction between phages and bacterial populations actively migrating through a porous environment. Bacterial migration is driven by chemotaxis up a self-organized nutrient gradient (13,14). Chemo-taxis is widespread in many natural habitats shared by bacteria and phages, including oceans (15), mucosal membranes in the gut (16,17), soil, and other environments reviewed in Ref. (18). Hence, it is likely that the dynamics elucidated here are relevant to a broad spectrum of real-world ecosystems.

Below we show that the interplay between chemotaxis and phage predation results in a rich spatio-temporal dynamics involving multiple rounds of phage infection separated by spatial distances measured in centimeters. The emergent property of this system is a propagating “bacterial” front composed of the fastest migrating bacteria, which expands symmetrically across an agar plate. This bacterial front is trailed by another, “phage”, front, at which the bacterial population collapses due to an exponentially increasing number of phages. The rate of migration of this phage front increases with the size of the initial phage population and the rate of infection until it overtakes the migrating bacterial front and forces its collapse. Remarkably, the phage front is not symmetric but is instead structured on a millimeter length scale. Such rough fronts emerge close to the transition region between phage- and bacteria-dominated regimes. We suggest that this roughness is due to the stochastic nature of multiple rounds of phage infections combined with positive feedback between the growth of the phage population and the local density of unconsumed nutrients on the plate.

## Results

To study phage-bacterial interactions in a spatially structured context we used rich medium (Lysogeny Broth (LB)) low viscosity (0.3% w/v) agar plates. When *Escherichia coli*(MG1655-motile) is inoculated at the center of these plates it rapidly consumes nutrients locally creating a gradient in the primary attractant (serine) that drives radially symmetric chemotactic migration of the population (~ 0.3 cmh^−1^) (see Ref. (14) describing a similar experiment, which does not involve phages). Here we inoculate the system with a mixture bacteria and phage (P1, virulent strain) at the center of the plate. Phage P1 requires calcium (CaCl_2_, ranging between 1mM and 10mM in our experiments) and magnesium (MgSO_4_, 5mM) to infect *E. coli*(19). This dependence allows us to modulate the adsorption rate of phage by varying calcium concentration in the plate (MgSO_4_ concentration is fixed at 5mM in all experiments shown here).

Fig. 1 shows the spatio-temporal dynamics of populations of bacteria (about 1 × 10^6^ cells/inoculum) and phages (about 1500 PFUs/inoculum) for 1mM CaCl_2_. At this relatively low calcium concentration, the phage adsorption rates are low. Fig. 1A shows an image of the agar plate taken 8 hours, 14 hours and 19 hours after the inoculation. The overall radial expansion of the bacterial population (referred to as bacterial front, bright outer ring of the colony) is taking place at a steady pace of 0.36 cm h^−1^. This rate of expansion is identical to bacteria migrating in the plates without phage present (14). The bacterial front is followed by a deepening and slowly widening black region in the center of the agar plate, whose outer boundary we refer to as the phage front. This central black region is caused by a collapse of the local bacterial population due to phage predation. Radial profiles of the bacterial density were measured by extracting and averaging pixel intensities in background-subtracted images (see Methods). Fig. 1B shows these radial density profiles of bacteria at the time points shown in Fig. 1A. Here the phage front is clearly visible as a sharp decline in image intensity near the center of the plate at later times (e.g. for orange trace showing radial profile at 19 hours, the radius of the phage front is around 2 cm, while that of the bacterial front is 6 cm). For the conditions shown in Fig. 1, the separation between the phage front and the bacterial front increases with time (compare red and orange traces, Fig. 1B). Thus under these conditions phages are not able to keep up with the expansion of the bacterial culture.

**Fig. 1.**
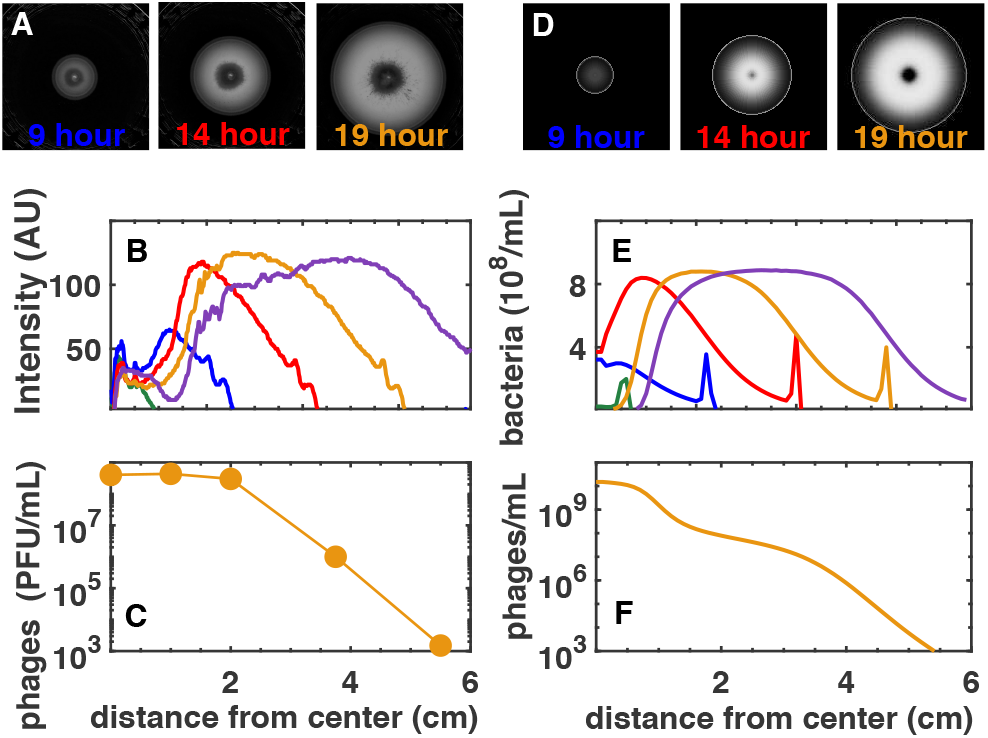
Spatio-temporal dynamics of bacteria and phage in a soft agar plate with 1mM concentration CaCl_2_. The center of the plate (radius 7.5 cm) was inoculated with approximately 1500 phages (PFUs/inoculum) and 1 × 10^6^ bacteria. (A) Images visualizing the spatial distribution of bacteria (light regions) on the plate at 8, 14 and 19 hours after inoculation. (B) The radial profile of the bacterial population density at 4 hours (green) 9 hours (blue), 14 hours (red), 19 hours (orange) and 24 hours (purple) after inoculation along a radial direction at angle 90° right from the vertical. (C) The radial profile of the phage population density measured at 19 hours. The phage density reaches a plateau around the same spatial position at which the bacterial population collapses (referred to as the phage front). (D-F) Are identical to (A-C) except show results of model simulations in which the phage latency time is inversely proportional to the bacterial growth rate (with offset), and phage burst size is proportional to the growth rate (See SI Appendix for details of the model).

To confirm that the collapse in bacterial densities at the center of the plate was in fact driven by phage predation and to measure phage densities in the expanding bacterial front, we sampled a plate after 19 h of expansion at several distances from the center and measured the phage density (Methods). Fig. 1C shows the phage population density as a function of radius where, as expected, we observe very high phage densities in regions where the image intensity is low. Surprisingly, even at such a late time-point, a small number phages are present at the outer edge of the expanding bacterial colony. The phage at the outer edge of the colony could not have moved there by diffusion alone (20). Instead, we speculate that the phage travel along with migrating bacteria during the latent infection period. However, given that the phage front is moving at a slower rate than the bacterial front, we expect that were the colony allowed to expand indefinitely, at some later time phages would completely disappear from the advancing bacterial front. At this point the population of chemotaxing bacteria would break free of their phage predators and be able to form a new phage-free colony.

To describe the spatio-temporal dynamics observed in Fig. 1A-C, we developed a deterministic ODE model of the phage-bacteria system. This model approximates the bacterial growth in LB medium using an (additive) Monod form for two food sources: one which acts as a chemoattractant (serine), and another representing all other nutrients present in the complex media, which do not drive chemotaxis. Furthermore, the model assumes that phage burst size decreases and the latency increases as bacteria growth slows when they use up their resources(21,22). In developing the model we found it necessary for phage adsorption rates and latency to depend on host physiology in order to reproduce the fact that the separation between phage and bacterial fronts increases with time. Without these physiological effects, the expansion of the phage-dominated central region proceeds at the same speed as the bacterial front migration rate. Batch culture experiments confirmed that this assumption of the model is reasonable for the phage P1-*E. coli* systems used here. Numerical simulations of the model are shown in Fig. 1D-F where we find that the model captures the core features of the data. A detailed description of the model is given in the Methods and SI Appendix.

Are bacteria always capable of outrunning their phage predators, or can phage populations slow down or even stop the bacterial front from advancing? To investigate this question we repeated our experiment for a broad range of parameters characterizing phage infectivity and initial population size. Fig. 2A shows snapshots of bacterial populations taken 19 h after inoculation, as a function of CaCl2 concentration on agar plates (x-axis) and the initial number of phages (PFUs) in the inoculum (y-axis). Higher concentrations of CaCl_2_ increase the adsorption coefficient of phages (19), thereby making it easier for them to spread. This is indeed seen as a systematic increase in the radius of the inner black region (phage front) as one moves from left to right in Fig. 2A, bottom row. Similarly, increasing the initial phage density speeds up the phage front migration even at fixed CaCl_2_ concentration (Fig. 2A, left column).

**Fig. 2.**
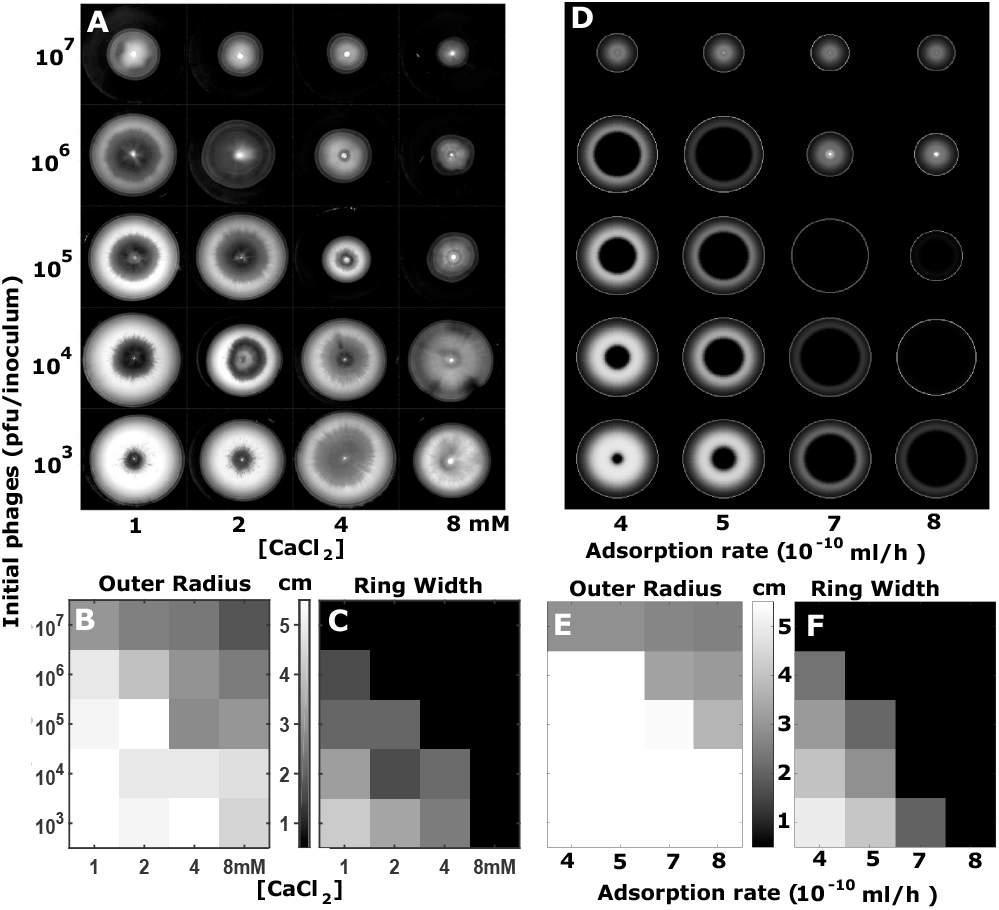
(A) Bacterial populations shown 19 h after inoculation with varying phage load quantified by the initial number of phages (PFUs) in the inoculum (y-axis), and CaCl2 concentrations in the agar. Note the logarithmic scale of both axes. The white intensity is proportional to the local density of bacteria with the inner black circle marking the area where bacterial population has collapsed due to phages. (B) The radial position of the expanding outer bacterial front at 19h. (C) The width of the region of high bacterial density between the outer bacterial front and the inner phage front. The position of the phage front is defined by the bacterial density dropping to 80% of its peak value. (D) Model simulations (see Methods for details) with resistant bacteria evolving at rate 10^−5^ per cell generation. (E-F) Bacterial front location and the width of the region of high bacterial density at 19 h in model simulations as in panel (D), but without resistant bacteria.

Remarkably, increasing both calcium concentrations and initial phage population size above a certain threshold results in a dramatically slowed bacterial front migration rates (upper left, upper right and lower right corners, Fig. 2A). Fig. 2A thus effectively acts as a phase diagram delineating conditions where bacteria win (lower left region) from conditions where phages win (upper right region).

Notice that in the region where phage population appears to overtake the bacteria, the collapse of bacterial populations is only temporary due to the ultimate growth of phage-resistant bacterial strains (23,24). For phage P1, these mutant bacteria naturally emerge at a rate of about 10^−5^ per generation (25). Clonal expansion started from single resistant mutants gives rise to radial sectors, reminiscent of spatial antibiotic resistance patterns (26) (Fig. 2A, 1 × 10^4^ phage, 8mM calcium). However, when the phage load is large enough at the beginning of the experiment, all susceptible bacteria are eliminated at the outset, resulting in delayed but spatially symmetric expansion of the resistant bacteria (Fig. 2A, upper right corner).

For large phage infectivity and abundances far above the transition line we observe the following interesting effect also recapitulated in our model (See the S1 Movie recorded for 10mM CaCl_2_ and 1.5 × 10^5^ PFUs/inoculum). For these parameters the advancing front of susceptible bacteria is completely stopped and decimated by phages around 12 hours after inoculation. At even later times, multiple clones of resistant bacteria emerge and spread away from the white ring marking the position of the stopped front of susceptible bacteria.

To quantify the migration of phage and bacterial fronts we measured their respective locations in space after 19 hours of incubation. Figs.2B and C show the position of the outer (bacterial) front and the width of the bright region between bacterial and phage fronts respectively measured 19 hours after inoculation (Fig. 2A). These figures illustrate the properties of the system on both sides of the “phage” transition: the bright area in the lower-left corner of the panel Fig. 2B shows that for low phage loads bacterial chemotaxis proceeds at a rate that is approximately independent of the phage load and calcium concentration. Time lapse imaging of expanding colonies at different CaCl2 concentrations showed that the migration rate of the bacterial front at low infectivity is identical to migration in the absence of phage. Further, these recordings show that the migration rate of the phage front increases with increasing CaCl2 concentration until the phage front ultimately overtakes the bacterial front at high phage loads (SI Appendix).

The progressive darkening of Fig. 2B in the upper-right corner quantifies the slowing of the migrating bacterial front by phages under increasing initial phage population sizes and infectivity. Above the “phage transition” where bacterial populations collapse and the migration rate of bacteria slows dramatically, we expected that the remaining bacterial front is driven by phage resistant strains. We tested this assumption by directly sampling bacterial populations from plates above and below the “phage transition” and assaying their ability to grow on lawns of phage. Indeed, above the transition the entire bacterial colony is comprised of resistant cells. In addition, at low infectivity and phage populations (1mM CaCl2, 1ğ55 × 10^5^ phage) only the bright center of the colony harbors phage resistant bacteria (SI appendix).

Fig. 2C illustrates how the inner (phage) front expands and gets closer and closer to the bacterial front until ultimately merging with it above the transition. This is quantified using the width of the ring where bacteria are abundant (the distance between positions of bacterial and phage fronts). In the black region in the upper-right corner of Fig. 2C phages spread as far as the bacterial front. Thus in this region no phage front is detected and the width is zero.

Overall Figs.2A-C show the existence of two distinct parameter regimes in which prey (bacteria) and predators (phages) dominate. As either the phage load (initial population) or phage adsorption rate (calcium concentration) increase, the system transitions from the regime dominated by bacterial prey to that dominated by phage predators.

Fig. 2D-F show that our deterministic ODE model reproduces the general properties of the measured phase diagram shown in Fig. 2A-C. The intensity profiles shown in Fig. 2D include the resistant bacteria, which dominate the bacterial populations in the upper right corner of the diagram but are not significant at or below the critical transition at the diagonal. Successful fits of experiments with our model require strong dependence of phage latency time (increase) and burst size (decrease) when bacteria slow down their growth due to nutrient limitation.

For conditions close to the transition region, where the phage front migration rate is only slightly slower than the bacterial front (e.g. 4mM CaCl_2_, 1 × 10^3^ phage, Fig. 2A) the phage front becomes rough, exhibiting a star-like spatial structure with centimeter-long phage-dominated dark regions (“death rays”) extending into lighter regions dominated by bacteria. This “death-star” phenomenon with large differences in bacterial densities spatially separated by as much as one millimeter suggests a role for contingencies in infection histories resulting in spatially structured bacterial populations.

To gain more insight into the microscopic phage-bacteria dynamics driving the transition between two regimes we studied a single plate in more detail. Fig. 3 shows two snapshots (19 and 24 hours after initiation) of a plate containing 5mM calcium and 4.7 × 10^3^ phage (PFUs) in the inoculum. These images reveal clear spatial structure in the phage front characterized by alternating black and white regions that are much more spatially extended than those shown in Fig. 1A. The speed of the radial expansion of black regions is faster than for 1mM calcium: along some angular directions the phage front advances at roughly the same speed as the bacterial front (compare the width of the light area near protruding black “death rays” at 19h vs 24h snapshots in Fig. 3AB).

**Fig. 3.**
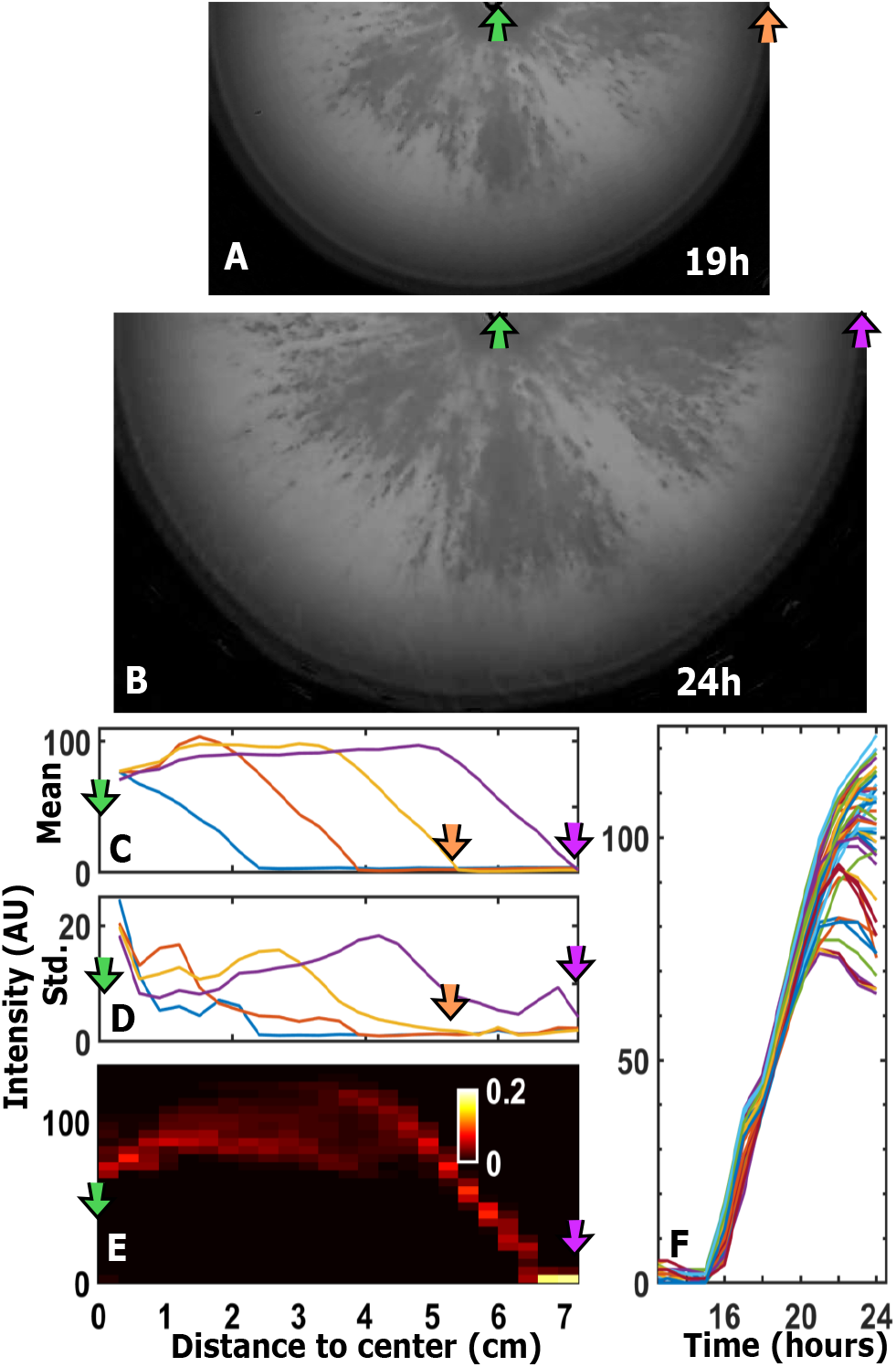
Bacterial population dynamics with 5mM CaCl_2_ and ~5000 phage (PFUs) in the inoculum quantified by pixel intensity in images with subtracted background. Half-plate images at (A) 19 hours and (B) 24 hours. (C,D) show average and standard deviation of pixel intensity as a function of the distance from the center at 9 hours (blue), 14 hours (red), 19 hours (orange) and 24 hours (purple). (E) 24 hour probability distribution (greyscale) of pixel intensity (y-axis). The distribution is normalized at each distance from the center (x-axis). (F) Time development of intensity for 50 different pixels located at different angles but the same distance (4 cm) from the the center of the colony.

To better quantify the spatial structure of phage “death rays” visible in Fig. 3AB, we computed statistics of image intensity (for background subtracted images) at fixed radii from the center of the plate (e.g. an annulus of pixels falling within a range of radii). Fig. 3C shows the azimuthally averaged image intensity as a function of radius at several times after the plate was initiated, while Fig. 3D shows the standard deviation of all pixel intensities at a fixed radius. Fig. 3D clearly shows that the variation in pixel intensities is maximal at intermediate radii (see peak around 4 cm for 24 hours purple trace). From Fig. 3C we infer that the width of the region where bacteria coexist with phages broadens with time much to the same extent as for much less aggressive phages shown in Fig. 1. On the other hand, the distance with the largest variation in bacterial abundances (the position of the standard deviation peak in Fig. 3D) moves forward at about the same speed as the advancing bacterial front.

To further elaborate on this in Fig. 3E we plot the normalized histogram of bacterial abundances (quantified by image intensity, y-axis) at 24 hours as a function of the distance from the center (x-axis). For distances around 4 cm (the position of the largest variation in Fig. 3D) one can see that the distribution of bacterial abundances is in fact bimodal, corresponding to white and black regions visible in Fig. 3AB.

Another way to quantify the bimodal nature of the phage dynamics near the transition is shown in Fig. 3F following the time course of bacterial abundance in 50 randomly selected pixels all located at 4 cm from the center. The bacterial front arrives to these locations nearly simultaneously around 15 hours after inoculation. Following near-synchronous increase during the next 5 hours, bacterial populations at different angular locations start to dramatically diverge from each other. Pixels which end the time series at lower intensities are inside “death rays” with high rate of phage-induced population decline. Conversely, pixels that remain at high intensities throughout correspond to white areas between death rays.

Therefore, for 5mM CaCl_2_ and 5000 phage (PFUs) in the inoculum, phages keep up with bacterial front expansion in some directions, but are left behind in others. This indicates that the local dynamics in our system has sufficient positive feedback to amplify initially small variations into macroscopic differences manifesting themselves at a scale of several centimeters.

## Discussion

We have shown how spatial structure in bacterial populations driven by chemotaxis interacts with phage population size and adsorption rate to determine the outcome of bacteria-phage interactions. Our study has three important results which should be considered in the context of phage-bacteria interactions in microbial communities from the human gut to the ocean.

First, we find that non-motile phage are able to “surf” expanding bacterial populations by repeatedly re-infecting cells at the expanding bacterial front (Fig. 1). This observation demonstrates that the latent period of phage infection serves as a mechanism by which phage are able to hitchhike with their hosts into new spatial niches. In fact, one can consider the spatial changes in bacterial and phage populations starting at the outer front and moving towards the center of the plate are roughly equivalent to temporal changes in a well-mixed phage-bacteria experiment. A handful of phages start the epidemics at the outer bacterial front. Bacteria behind the front do not chemotax due to a lack of nutrient gradients. So, here both phage and bacterial populations for the most part grow locally. Due to the constant speed of expansion of the outer front, the time profile of this local growth is approximately repeated as a spatial profile: the larger is the distance to the the bacterial front, the longer this population had to grow since it was “inoculated” with a mixture of bacteria and phages by the passing bacterial front. Hence, the phage population is expected to grow exponentially as one moves from the front towards the center of the plate (see Fig. 1C) until it reaches the level necessary to decimate the bacterial population at the phage front (see Fig. 1B).

Second, Fig. 2 shows how the outcome of phage bacteria interactions depend on population size and adsorption rates. A “phage transition” defines a boundary between two regions of parameter space, one where phage-susceptible bacteria escape phage infection, and another one, where the bacteria are forced to evolve phage resistance to survive. This observation has important implications for understanding phage-bacteria interactions in complex environmental contexts such as lakes or the particle-filled water columns in the ocean, where bacterial growth is variable due to transient chemotaxis driven exploitation of patchy resources (15). In particular, for phage to dominate a migrating and growing bacterial population requires high initial population densities or high adsorption rates. In situations where these conditions are not met, phages remain able to attack smaller localized bacterial populations, which limits their overall fecundity and possibly their impact on bacterial genome evolution via transduction (27).

A key feature responsible for the spatial structure in phage-bacteria populations is the strong dependency of phage infection on host physiology. The increasing burst size and decreasing latency of phage infections with host growth rate interacts with the self-organizing nutrient gradients in migrating bacterial populations to give rise to regions of space where phage are excluded. Previous studies suggest this dependence on host physiology arises from limits on the rate of synthesis of new phages and on the size of host cells prior to lysis(22).

When either phage infectivity or the initial density of phage in the inoculum are high, the phage population at the moving front amplifies exponentially, ultimately leading to a collapse of the bacterial front. Our model correctly reproduces this transition as a function of the phage in the inoculum, see Fig. 2D. Above the transition, the food is not depleted substantially by susceptible bacteria prior to collapse so that phage-resistant bacteria can grow and reach high densities. As the initial fraction of resistant bacteria is small, their final growth to a propagating front of high density is delayed. A similar process happens in batch culture when a large phage population is initiated with a small bacterial population in well-mixed conditions (Fig.S2A). At high CaCl_2_ concentrations and initial phage populations that are close to but above the transition line, Fig. 2A shows “clonal fans” of resistant mutants reminiscent of fans of antibiotic-resistant clones in Ref. (26). Some of the individual fans can be traced back to the original inoculum, while others arise from different points at the circle where the first wave of susceptible bacteria has collapsed under phage pressure. The collapse of the susceptible population presumably left much of the nutrients untouched resulting in subsequent growth of the resistant population. You can see the emergence of resistant “fans” in Supplementary Movie 1 showing the dynamics with 10mM CaCl_2_ and 1.5 × 10^5^ PFUs/inoculum.

Third, a striking result of our study is the extent of contingency manifesting itself as the “death star” (Fig. 3AB), where centimeter-wide spatially extended regions have very different fates. A full accounting of this phenomenon awaits future work, however, dramatic differences in bacterial populations of billion cells are amplified from relatively small fluctuations in the inoculum with a large number of phages (between 10^3^ and 10^7^ in our experiments).

At first glance, the alternating black and white regions in a death star, observed for high calcium concentrations close to the transition (see Fig. 3A), are reminiscent of the emerging sectors in microbial range expansion experiments in Ref. (28). This analogy can be partially justified by criticality in the sense that phages are neither winning nor loosing the race with bacteria. Hence the “expansion fitnesses” of dark and light regions are approximately equal to each other. The same is true of fluorescently labeled identical microbial strains in Ref. (28). One must note, that in our experiments bacteria migrate in the radial direction. This gives rise to much straighter boundaries between sectors than a random walk reported in Ref. (28). Another distinguishing property is that sectors in our experiments are *not caused by mutant strains* of either bacteria or phages. Indeed, the resistant bacteria typically enter the game at a later time and have very different fan-like spatial abundance profiles similar to those reported in Ref. (26). Instead, the sectors in our system are the emergent property of the spatio-temporal population dynamics of phages and bacteria.

To better understand these spatial fluctuations will require a combination of: 1) detailed understanding of spatially-localized phage fluctuations within the inoculum before and just after the start of migration; 2) the mechanism of positive feedback during multiple rounds of infection within the advancing front.

One possible mechanism for the positive feedback is that phage populations, which initially succeed in killing the bacteria thereby prevent these bacteria from consuming the nutrients that the phages need for subsequent proliferation on growing bacteria. Ref. (29) used the Lotka-Volterra framework to demonstrate that reduced adsorption rate as bacteria approach stationary state can provide sufficient feedback to give rise to bi-stability. Our deterministic model similarly takes nutrient dependence into account by changing both burst size and latency time.

A close inspection of the light regions in Fig. 3AB shows that the black areas slowly “eat” into them over time. This could be tentatively explained by food diffusion. In conclusion, positive feedback caused by details of phage physiology may explain how regions with high bacterial density can survive in the presence of large density of phages in the neighborhood. However, it leaves unexplained where the dark regions are coming from when the number of phage in the inoculum is relatively large.

To explain this we conjecture that each black ray is associated with an early lysis events of a single infected founder bacteria at the outer edge of the initial inoculum. Since abundant nutrients at the bacterial front favor phage proliferation, an early burst of phages may establish a small droplet of infected bacteria migrate together at the front. Phages left behind by this droplet may subsequently infect and would ultimately cause local collapse of host population along newly formed black rays of the death star. In this case the width of the “death rays” would be determined by the bacterial diffusion constant suggesting a future statistical analysis of the imaging data presented here.

In all cases, actively moving hosts impose a new set of selection pressures on their viral predators. For example, we expect that phages with infection dynamics inhibiting chemotaxis would be at a disadvantage. Indeed, such phages would not be able to hitchhike inside their bacterial hosts and thereby would be left behind. Chemotaxing bacteria thereby add an additional constraint of “gentleness” selecting against phages with extremely aggressive infection cycles.

## Materials and Methods

### Experimental methods

All experiments were performed with LB rich medium (10 gL^−1^ Tryptone, 5gL^−1^ yeast extract, 5gL^−1^ NaCl). To create a swarm plate LB medium was autoclaved with 0ğ3 %w/v agar after autoclaving 5mM MgSO_4_ and an appropriate CaCl2 concentration were added. Prior to cooling 100 mL of medium was measured in a sterile graduated cylinder and poured into a 15 cm diameter sterile plate. The plate was allowed to solidify at room temperature for 3 hours. Plates were then transferred to a walk in incubator 30°C (Darwin Chambers) and allowed to reach thermal equilibrium for no less than 24 h prior to inoculation. All experiments reported here were performed at 30 °C.

To initiate an experiment an overnight culture of *E. coli*(MG1655, motile, Coli Genetic Stock Center #6300) was initiated in LB from frozen glycerol stocks. 10 μL of saturated overnight culture (≈ 1 × 10^6^ cells) was taken from this culture and mixed with 10 μL phage P1 lysate and vortexed throughly before being inoculated into the center of the swarm plate. P1 lysates were titered by standard serial dilution and plating on top agar and counting plaques. Lysate titers were ~1 × 10^10^PFUs/mL. Changing the initial phage population was accomplished by diluting the P1 lysate into LB medium prior to mixing with bacteria. Phage lysates were not used for more than a year and each new phage lysate that was made was titered by serial dilution and plating.

Plates were imaged using computer controlled Logitech webcams housed in light tight boxes with illumination provided by LED strips which encircled the plates and were turned on for one second per image acquisition (14). Imaging occurred in one of two ways: either plates were left in light tight boxes and images were acquired once every two to four minutes or plates were housed in the incubator but were manually transferred to the imaging apparatus to acquire images at fixed points in time since inoculation (Fig. 2). All image analysis was performed with custom written Matlab scripts.

Plate reader experiments were performed in 48 well plates using a BMG Labtech Clariostar plate reader equipped with an incubator maintained at 30 °C. Media volumes in each well were 0.75mL. The plate reader made automated optical density measurements on 5 minute intervals with orbital shaking between measurements.

### Model

To interpret the experimental observations we used a continuous population dynamics model in which local (1mm×1mm on the plate) abundances of both bacteria and phages are described by a set of deterministic differential equations in space and time. The growth medium is modeled as a (10%-90%) mixture of two nutrients, with the less abundant nutrient also acting as a chemoattractant. The depletion of this chemoattractant causes bacteria to chemotax away from the center of the plate. The subsequent bacterial growth behind the front is primarily due to the second nutrient. Phage infections are described using traditional Lotka-Volterra equations (1,2). Importantly, in order to reproduce the experimental results we had to add to our model a dramatic reduction of phages’ burst size and extension of their latency time as nutrients get depleted and local bacterial populations approach stationary state. A very slow diffusion of phages (1 mm over 24 hours) is ignored in our model. More detailed description of our model is provided in the SI Appendix.

## ACKNOWLEDGMENTS

This research has received funding from the European Research Council under the European Union’s Seventh Framework Programme (FP/2007 2013)/ERC Grant Agreement n. 740704. D.P. and S.K. acknowledge support from the National Science Foundation Physics Frontiers Center Program (PHY 0822613 and PHY 1430124). This work was performed in part at the Aspen Center for Physics, which is supported by National Science Foundation grant PHY 1607611.

## Supplementary Appendix

### A. Spatial structure of bacterial populations resistant to phage

Our model assumes that above the “phage transition” the migrating bacterial front is comprised solely of bacteria resistant to phage infection. Conversely, in conditions well below the transition, we expect that only a small population of cells at the center of the plate are resistant to phage infection. To test this assumption explicitly we ran two separate plates with 1.55 × 10^5^ phage in the inoculum and with 1mM (below the transition) and 8mM CaCl_2_ (above the transition) calcium concentrations. After a fixed period of time we sampled bacteria at different distances from the center of the plate and tested them for susceptibility to phage infection by streaking cells on a hard agar plate where a strip of the plate was inoculated with phage. We expect resistant bacteria to grow in the region of the plate where phage are present and susceptible bacteria to be killed in this region. As expected, we find that the outermost bacterial front is susceptible to phage infection in the 1mM condition (Fig. S1, left column, samples A,B,C, F,G), while only bacteria at the very center of the plate are resistant to infection (Fig. S1, samples D, E). Conversely, all populations sampled from an 8mM plate were found to be resistant to phage infection (Fig. S1, samples H-N). Note that the trailing off of bacterial densities in samples H-N can be attributed to the substantially lower density of bacteria in the 8mM condition after 19 hours of growth.

**Fig. S1.**
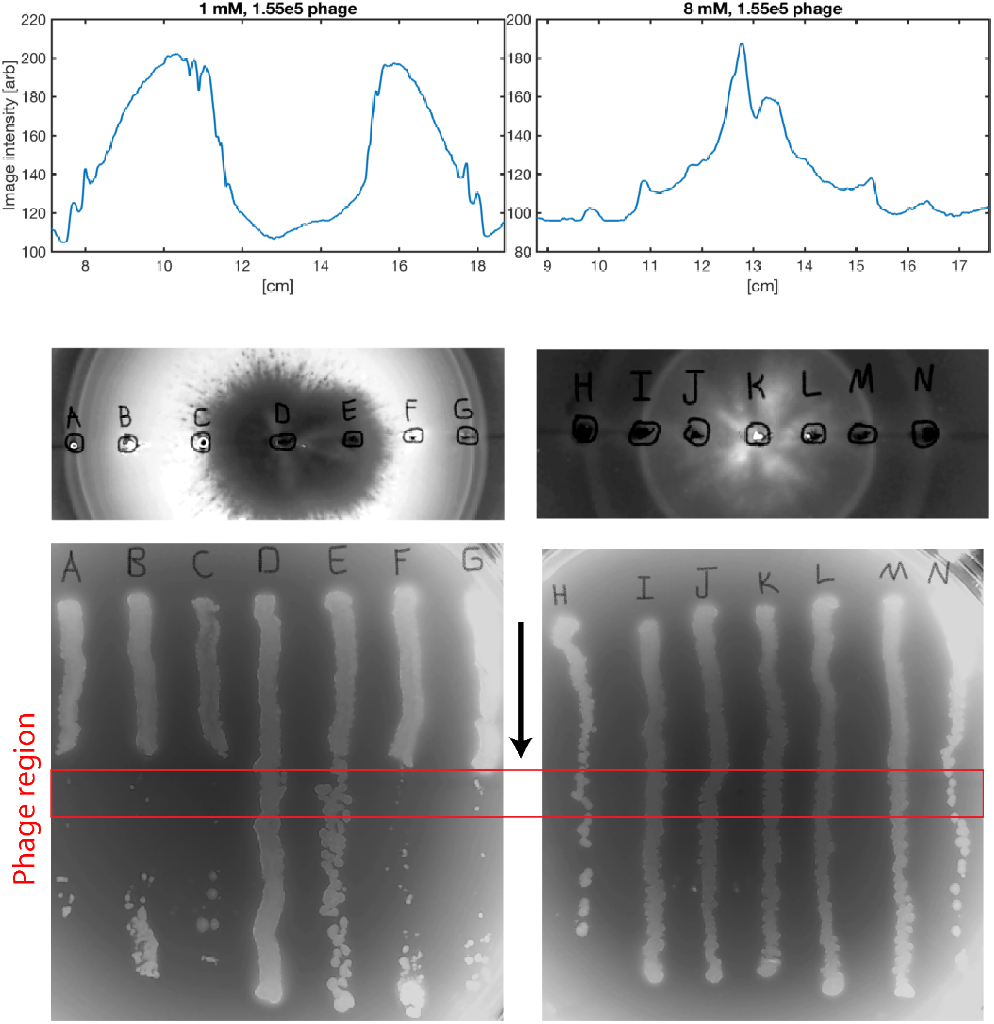
Outer bacterial fronts are susceptible (resistant) to phage below (above) the transition. Two plates in conditions below (left column, 1mM CaCl_2_, 1.55 × 10^5^ phage) and above (right column, 8mM CaCl_2_, 1.55 × 10^5^ phage) taken after 19 hours of growth and migration. Top row shows image intensity profiles through the center of the colonies. Middle row shows images of plates with points labeled by the letters. Images in the middle row are aligned with intensity profiles in the top row. Samples from each point were streaked on hard agar plates which had a strip of agar inoculated with high densities of phage P1 (red box, bottom row). Cells were streaked in the direction shown by the black arrow (middle).

#### Growth rate dependent infectivity

Our model assumes that phage are less effective in infecting bacteria as the bacterial growth rate slows. In order to test this assumption we performed a batch culture experiment where well-mixed bacterial cultures were infected with high doses of phage (3.6 × 10^8^ PFU/mL) at different points during the growth curve. To accomplish this, an overnight culture of bacteria in LB broth was diluted 100-fold into a 48-well microtiter plate. The plate was grown at 30 °C with orbital shaking in the plate reader with OD600 measurements taken every 5 minutes. Every hour the plate was removed from the plate reader and a subset of wells was infected. Indeed, for wells infected very early in the growth process we observe a long lag before a resistant population arises Fig.S2A. For cultures infected 4 hours after initiation an initial rise of presumably susceptible bacteria is killed by phage before a resistant population is established at long times (Fig.S2B). However, for populations that are infected 10 hours after initiation, when the bacterial growth rate has already slowed due to nutrient depletion, no decline in bacterial densities is observed (Fig.S2D). This observation strongly supports our claim that the ability of phage to infect bacteria declines with decreasing bacterial growth rate. Interestingly, at intermediate infection times we find that the final bacterial density is not reproducible, with some cultures remaining at high bacterial densities over long times, while others are reduced to low densities by phage killing (Fig.S2C). This apparent stochasticity likely results from well-to-well variation in the bacterial growth rate (and therefore phage infectivity).

**Fig. S2.**
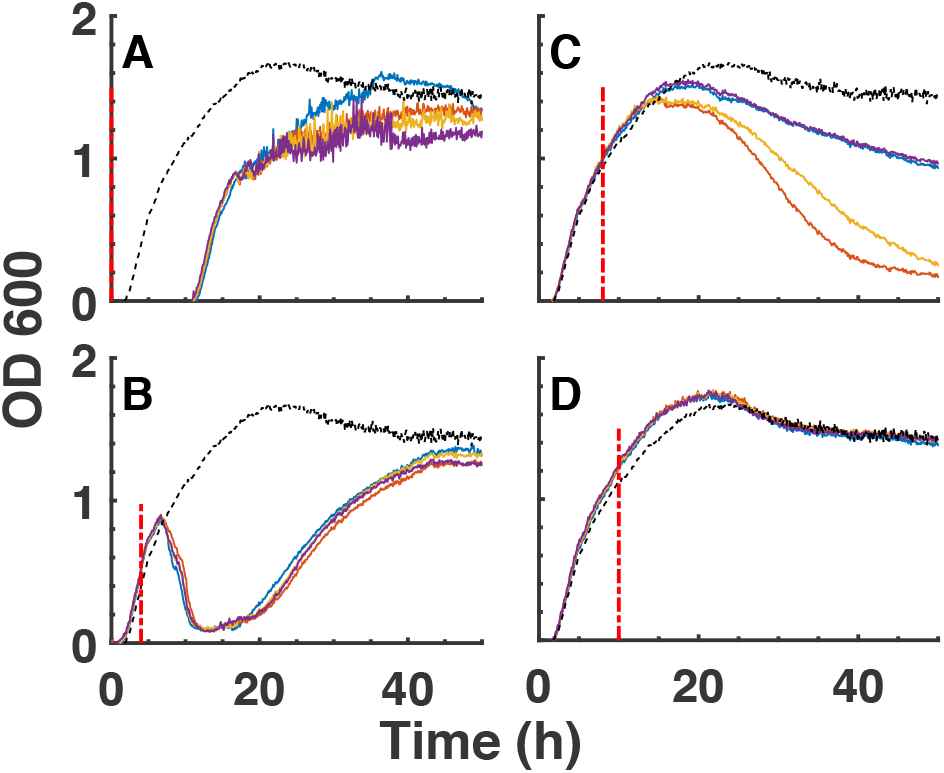
Batch culture development, starting with 2 × 10^7^ bacteria in 0.75mL LB medium with 10mM CaCl_2_. Panels correspond to different experiments where 2.6 × 10^8^ PFUs were added to the system at different time points marked with red vertical lines. Each panel show 4 independent replicates. A) Phages added at 0 hours. B) Phages added at 4 hours. C) Phages added at 8 hours. D) Phages added at 10 hours. In all cases the dashed curves mark the average dynamics of 4 replicates without phages.

#### Detailed description of computational model

Our model simulates the average population dynamics of bacteria and phages in our system on a lattice made out of 150 × 150 sites, each representing a 1mm× 1mm square on an agar plate used in our experiments. Since our model uses deterministic differential equations and the solution is naturally radially-symmetric, we simulated the radially-symmetric 1D version of equations below with appropriate expressions for gradients and diffusion terms. The dynamics of the phage population (P), different bacterial subpopulations (*B*, *I_i_*, *B_r_*), and concentrations of the chemoattractant/nutrient 1 (a) and the nutrient 2 (c) are described by the following equations

**Fig. S3.**
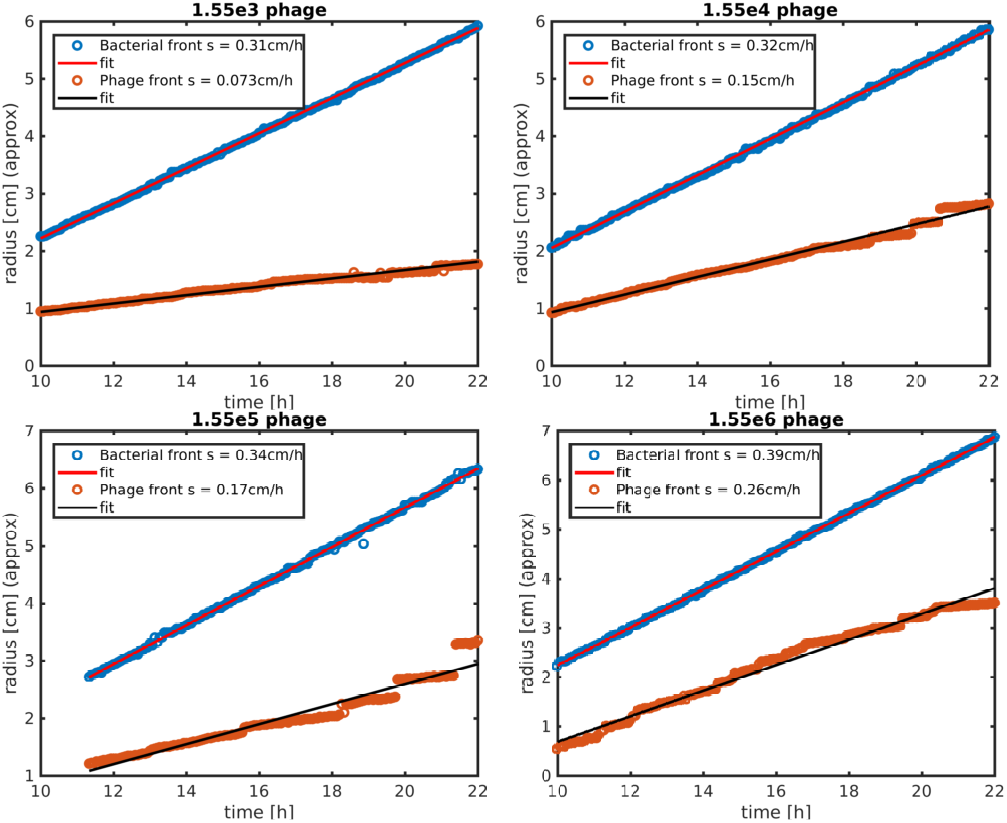
Migration rates at 1mM CaCl_2_. Migration rates of bacterial fronts and phage fronts for four plates at different initial phage population sizes corresponding to the left column of Fig. 2A. The front locations in time are measured from radial image intensity profiles for background subtracted images (taken at 90° from vertical) and averaged over a swath 8 pixels wide (e.g. Fig. 1A). The bacterial front location is determined by detecting a peak in the image intensity at the outer edge of the colony. The phage front location is determined to be the point along the radial intensity profile where the intensity falls to 80 % of its maximal value. Results are not sensitive to this threshold. Errors in the regression of front location in time are of order 0.01 cm h^−1^.

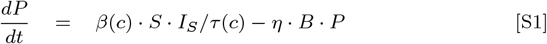

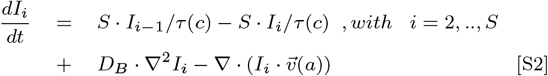

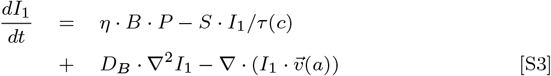

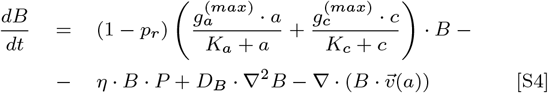

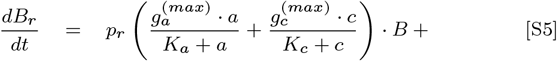

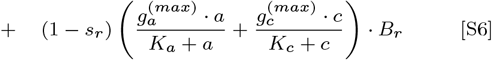

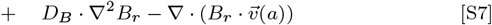

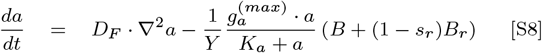

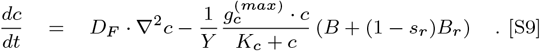

Here *F* = *a* + *c* represents all nutrients in the medium. Nutrients are assigned at an initial fixed concentration at each lattice point and are subsequently depleted due to bacterial replication. Initial concentration of *a* and *c* is *a*_0_ = 2*mM* and *c*_0_ = 18*mM* respectively. The bacterial growth is the sum of two Monod growth curves. The first contribution comes from growth on chemoattractant, *a*, doubling as a food source. This resource allows for somewhat faster bacterial growth rate. However, it accounts for only 10% of the total nutrient concentration *F*. The other nutrient, c, accounts for the remaining 90% of *F* but allows for somewhat slower maximal growth rate and dramatically larger Monod concentration *K_a_* (see the Table 1 for details). All these parameters were adjusted to best reproduce the growth curves of bacteria as they approach the stationary state. Both types of food sources are allowed to diffuse with a diffusion constant of *D_F_* = 3.0mm^2^/h (14).

**Table 1.**
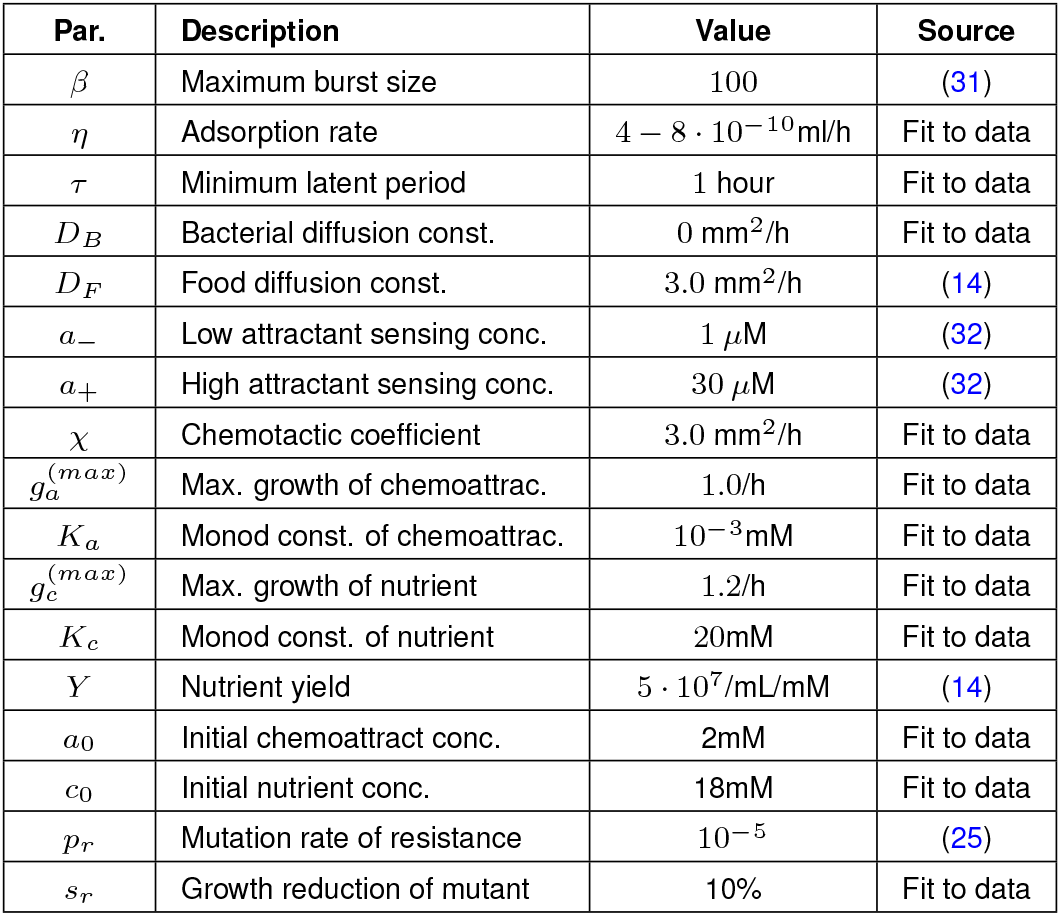
Model parameters used in Figs. 1 and 2.

The chemoattractant-dependent velocity 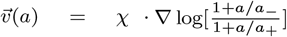 (30); models bacterial chemotaxis up the gradient of *a* with the adaptation range between *a*_−_ and *a*_+_. In addition, bacteria (which are actively moving in our case) have a diffusion constant *D_B_* ~ 6mm^2^/h (14), but we obtained somewhat better fit to our data by using *D_B_* = 0. In order to generate the bacterial fronts above the transition region in Fig. 2A, we introduced phage-resistant bacteria, *B_r_*, that grow *s_r_* = 10% slower than the susceptible bacteria, *B*. The generation of a phage-resistant mutant is assumed to be a rare event happening with probability *p_r_* = 10^−5^ per replication.

The parameters describing the dynamics of phage infections are: *η* - the constant adsorption rate, *β*(*c*) - the burst size, and *τ*(*c*) - the latency time. The latency is modeled by introducing *S* = 10 auxiliary intermediate infection states of bacteria, *I*_1_, *I*_2_, … to *I*_1_ taking place between the adsorption event and the production of phage progeny, each with average lifetime of *τ*(*c*)/*S* (10). By changing the number of intermediate steps *S*, one could explore effect of variance in latency time of individual bacteria. Indeed, for *S* = 1 the latency time is exponentially distributed and hence has standard deviation equal to the average. For other values of *S*, the overall latency time from infection to lysis would follow Gamma (or Erland) distribution with the same average *τ*(*c*) but with std/mean equal to 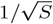. The intermediate steps are needed to model a lower bound in the latency time, characteristic of real phage infections. We verified that our results are not sensitive to *S* as long as *S* ≥ 10. At all stages, infected bacteria in states *I_i_* are assumed to move by diffusion and chemotaxis exactly as non-infected bacteria. However, they do not grow or consume nutrients and cannot be infected by phages for a second time.

In the absence of latency, phages would be able to move only by diffusion. For a realistic diffusion constant for phage P1, they would be able to spread by less than 1mm (12) within 24 hours duration of our experiment. Since the phage diffusion is so slow we set it to zero in our simulations. An important consequence of very low phage diffusion is that, both in the experiments as well as in our model, phages could move long distance (up to 7.5cm) only by hitching a consecutive series of “rides” inside infected bacteria.

As seen from Fig.S2CD, it is harder for phages to kill bacteria when growth conditions approach the stationary state. In order to reproduce the experiments from Figs. 1,2, we need to incorporate such attenuation of phage infections into our model. When nutrient gets used up, we assume that the burst size *β*(*c*) decreases and the latency time *τ*(*c*) increases as 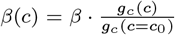 and 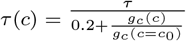. Here 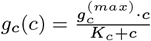 is the growth rate contributed by the nutrient *c* and 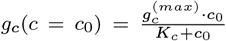 is initial specific growth rate on this nutrient with initial concentration *c*_0_ = 18*mM*. Parameters can be found in Table 1.

## Notes

Authors declare no conflicts of interest.

